# Alpha-tocopherol conjugated DNA tetrahedron with enhanced cellular uptake and selective cytotoxicity for cancer therapeutics

**DOI:** 10.1101/2025.05.12.653403

**Authors:** P Chithra, Payal Vaswani, Dhiraj Bhatia

**Affiliations:** Department of Biological Sciences and Engineering, Indian Institute of Technology, Gandhinagar, Palaj

**Keywords:** DNA tetrahedron, alpha-tocopherol succinate, selective cytotoxicity, covalent conjugation

## Abstract

One of the most fatal diseases in the world, cancer, lacks proper therapies that are toxic to cancer cells and specifically kill them. A newly emerging field, DNA nanotechnology facilitates the design of programmable, biocompatible DNA-based nanostructures, with applications spanning drug delivery, biosensing, and a number of applications in biomedical and therapeutics. However, the negatively charged outer leaflet of the plasma membrane poses challenges for the uptake of negatively charged DNA nanostructures. Strategies such as functionalizing DNA nanostructures with cationic lipids have been attempted, but these approaches have yielded conflicting results and certain limitations including stability and ambiguity of lipid functionalisation. Additionally, drug delivery using DNA tetrahedron (TD) and other conventional therapies has shown off-target effects due to the non-specificity of the drug. To address these challenges, this study utilizes a hydrophobic molecule, alpha-tocopherol succinate (AT), known for its selective cytotoxicity towards malignant cells over normal cells at appropriate concentrations. Covalently conjugating AT with TD preserved its selective toxicity property and enhance the cellular internalisation of DNA tetrahedron in specific cell lines. ROS generation was increased and led to apoptosis in malignant cell lines specifically. This suggests the development of a novel system with specific cytotoxicity towards cancer cells with increased uptake.

## 1. INTRODUCTION

Cancer remains one of the primary causes of mortality worldwide, responsible for nearly 10 million deaths in 2020 [1]. Projections from the Global Cancer Observatory suggest that the number of new cancer cases in India will increase from 1.4 million to 1.75 million by 2030 which is a 24% rise [2]. Of the 20 million global cancer cases recorded in 2020, breast cancer accounted for 2.3 million diagnoses and resulted in approximately 685,000 deaths [3]. Breast cancer is the most frequently diagnosed cancer among women globally, and in India, its incidence is rising rapidly, particularly among younger women [4]. Delayed diagnosis, limited awareness, inadequate treatment access, and poor follow-up care contribute significantly to the low survival rates of breast cancer patients in India [5], [6].

Conventional treatment strategies including radiation therapy, chemotherapy, peptide-based drugs suffer from drawbacks like damage to surrounding tissues [7], off target effects and systemic toxicity [7], [8], poor bioavailability and retention time [9], [10] respectively hinders their ability to stand out in the cancer treatment market. Targeted therapies, such as immunotherapies including monoclonal antibody production, immune checkpoint inhibitors, CAR-T cell therapy, and therapeutic vaccines offer high specificity, rapid response, and effective targeting of diseased cells [10]. For instance, trastuzumab (Herceptin) binds specifically to HER2 receptors on metastatic HER2-positive breast cancer cells, effectively slowing disease progression [11]. Despite their therapeutic promise, these treatments face several limitations, including high production costs, inherent immunogenicity, short in vivo half-lives, and poor tissue penetration [10]. While antibody-drug conjugates (ADCs) offer advantages such as targeted cytotoxicity and a well-characterized mechanism of action, they also present challenges [10], [12]. These include their relatively large molecular size, limited internalization efficiency, time-intensive processes for antigen identification and validation across different cancers, and variability in antigen expression among tumor cells all of which hinder efficient commercialization and broader clinical application [13], [14], [15]. These limitations highlight the urgent need for delivery systems that are inherently biocompatible, selectively cytotoxic, and structurally programmable.

Nadrian Seeman pioneered the use of synthetic oligonucleotides to build branched DNA motifs, using Watson-Crick base pairing laying the foundation for structural DNA nanotechnology [16]. In 1982, he showed that sequence-specific hybridization and sticky-ended ligation could form stable 2D and 3D DNA frameworks [17]. The field advanced in 2006, with Paul Rothemund’s development of DNA origami, where long DNA strands are folded into defined shapes using short staples [18], [19]. This enabled the creation of complex nanostructures used in drug delivery, biosensing, and nanoelectronics [20], [21], [22], [23], [24], [25]. Despite their promise, challenges like enzymatic degradation and scalability persist, driving innovations such as enzymatic amplification and chemical modifications to improve stability and function [26].

DNA tetrahedrons (TDs) have gained prominence among DNA nanostructures due to their structural stability, programmability, and biocompatibility [27], [28], [29]. DNA tetrahedrons (TDs) have been used to deliver chemotherapeutic drugs like doxorubicin and cisplatin through intercalation and electrostatic interactions [30], [31]. However, non-covalent loading poses challenges such as instability under physiological conditions and non-specific drug release due to interference from cellular components. These factors reduce drug availability at target sites and increase off-target toxicity [32]. Additionally, TD uptake is limited by the negatively charged DNA backbone and attempts to improve this with cationic lipid functionalisation have shown variable results [33], [34]. Covalently attaching a hydrophobic therapeutic agent to TDs offers a promising solution by enhancing stability, minimizing off-target effects, and improving delivery efficiency.

Alpha-tocopherol succinate (AT), a redox-silent vitamin E derivative, complements DNA-based delivery systems with its selective pro-apoptotic effect on cancer cells [35]. Unlike traditional antioxidants, AT targets mitochondrial complex II, disrupting electron flow and increasing ROS, which triggers cytochrome c release, caspase activation, and apoptosis [36], [37]. Since it doesn’t kill normal cells, it poses lower systemic toxicity and thus plays a dual role as pro-oxidant in tumor cells and antioxidant in normal cells [38], [39]. However, AT faces challenges such as esterase sensitivity, poor solubility, low bioavailability, reduced activation at high doses, and burst release in conventional delivery systems, all of which limit its efficacy [39]. Nanotechnology-based delivery can overcome these issues by improving stability, bioavailability, controlled release, and enhancing tumor targeting via the EPR effect [40].

In this study, we aimed to address a major challenge in cancer therapeutics i.e. non-specific cytotoxicity. Covalent linkage of AT and TD may overcome individual limitations, enabling improved stability, uptake, and cancer-specific cytotoxicity. We hypothesized that by conjugating or loading a hydrophobic molecule that selectively targets cancer cells, along with TD, we could enhance the overall uptake of the therapeutic system and reduce off-target effects. To test this hypothesis, we conjugated alpha-tocopherol succinate with a DNA tetrahedron and carried out invitro studies to prove the ability of the system to confer cytotoxicity only to malignant cells with higher cellular uptake.

## 2. RESULTS AND DISCUSSION

### 2.1 AT coupling with amino-modified M1 using HOBt-EDC chemistry

To facilitate coupling through HOBt-EDC chemistry, the M1 oligonucleotide was modified with an amino group in ts 5’ end and it was purchased from Sigma. Since the reaction should take place in a moisture-free environment for the formation of esters, the solvent and reaction conditions were optimized. EDC reacts with the carboxylic group of AT forming a O-acyclisourea intermediate. This intermediate is prone to hydrolysis or rearrangement to an inactive N-acylurea. HOBt reacts with O-acylisourea to obtain a stable HOBt-active ester which is resistant to hydrolysis and less electrophilic. The 5’ amino group in the modified M1 strand acts as a nucleophile which attacks the electrophilic carbonyl carbon of the HOBt-active ester. Consequently, amide bond is formed which covalently links AT with amino-modified M1 (Figure 1 a). Gel electrophoresis and UV-Visible spectrophotometry were used to confirm the coupling of AT with amino-modified M1 strand.

**Figure 1.**
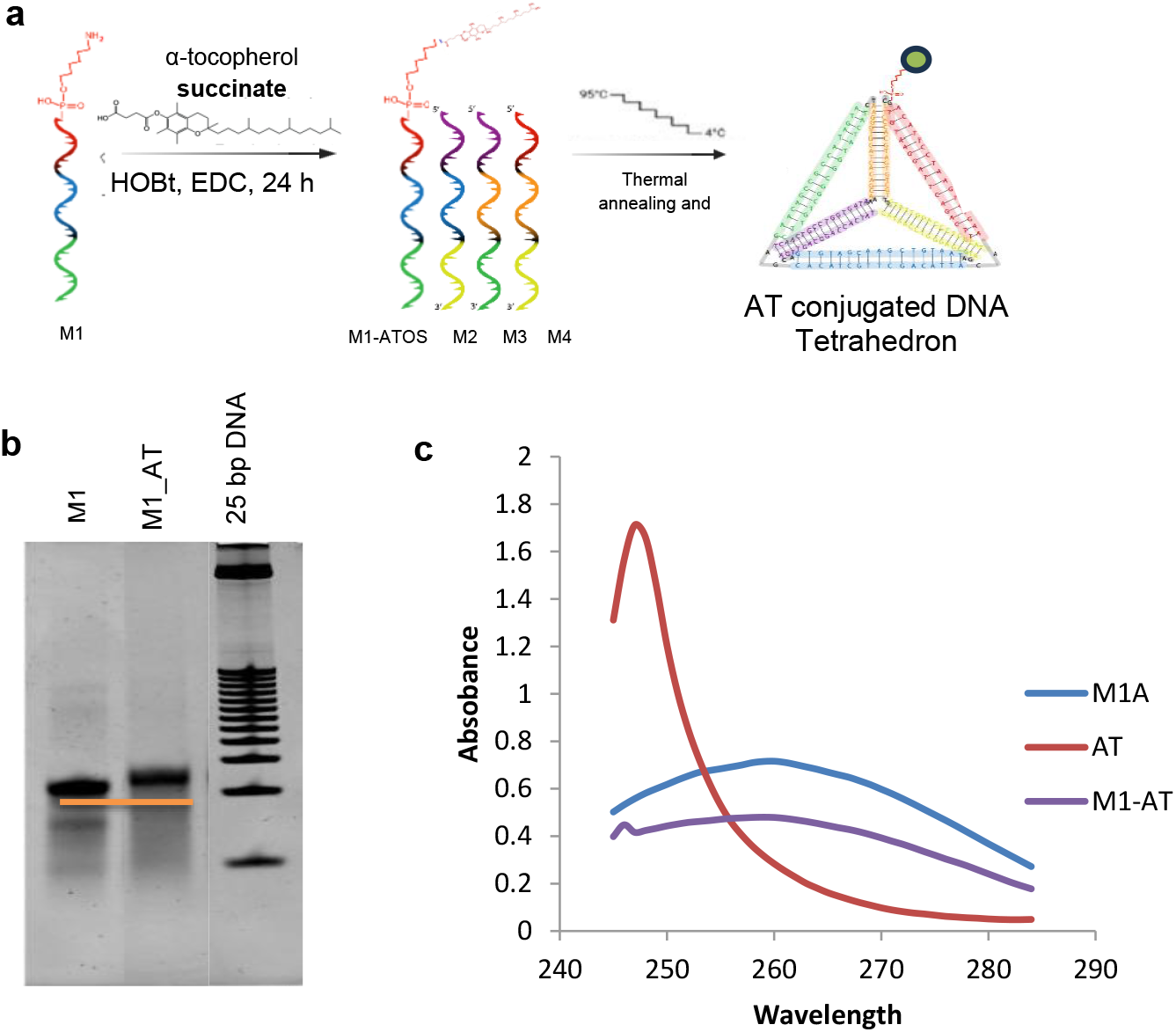
Coupling of AT with amino-modified M1 strand & characterisation. (a) Schematic Representation of AT coupling and TD_AT synthesis (b) 10% native PAGE showing the shift in band due to the occurrence of coupling. (c) UV-Visible spectrophotometry showing the characteristic peak of both AT (around 250 nm) and DNA (260 nm) in M1_AT.

In Figure 1b, due to increase in the molecular weight by 512.78 Daltons, there is a slight shift in the band was observed in AT conjugated M1 (M1_AT – 17571.78 Daltons) when compared to the non-conjugated amino modified M1 strand (M1 – 17059 Daltons). This is a preliminary confirmation of successful coupling of AT with M1. In Figure 2.1 c, M1_AT curve shows a significant dual absorbance at ∼250 nm (characteristic of AT) and at ~260 nm (characteristic of DNA – M1) due to the presence of both AT and M1 which suggests that AT is coupled with M1. This further validates the results obtained from gel electrophoresis studies.

**Figure 2.**
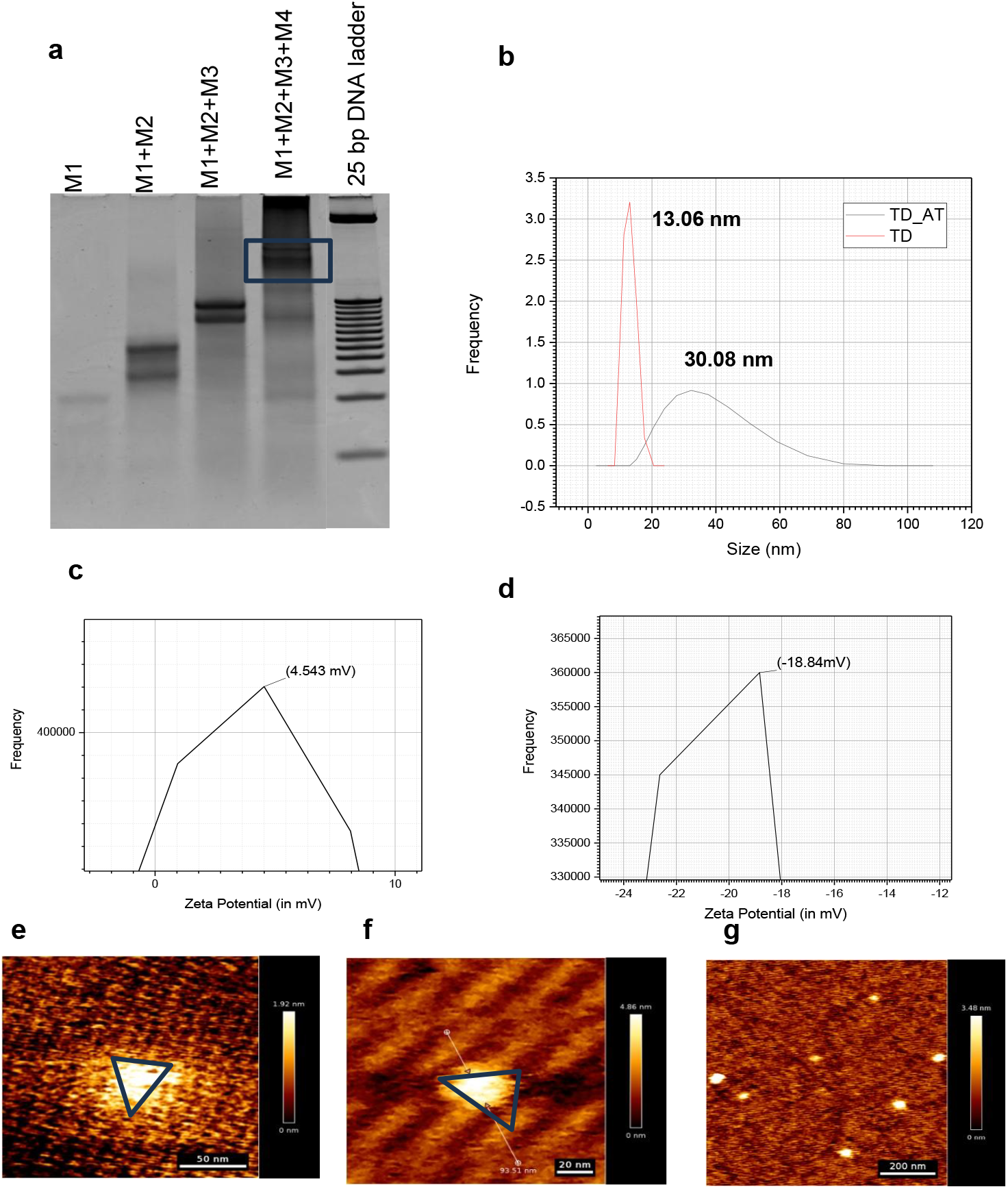
Synthesis of TD_AT & characterisation. (a) 10% native PAGE of EMSA showing the retardation in mobility due to the formation of TD_AT (b) Hydrodynamic size of TD and TD_AT measured using DLS. Zeta potential of (c) TD and (d) TD_AT. Atomic Force Microscopy images showing the formation of (e) TD & (f) & (g). TD_AT Scale bar represents 50 nm for (e), 20 nm for (f) and 200 nm for (g)

### 2.2 TD_AT synthesis & characterisation

The AT coupled oligonucleotide (M1_AT) was then employed to synthesize DNA tetrahedron using the thermal annealing protocol in a one-pot synthesis method along with three other oligonucleotides (M2, M3, M4). Confirmation of which was carried out using EMSA. EMSA depicted a ladder-like pattern as the number of oligonucleotides in the reaction increased. The retardation in the mobility of the bands were also observed which indicates the formation of higher-order DNA nanostructure (TD_AT) being formed as the number of primer increases (Figure 2 a). The average hydrodynamic size and the zeta potential of the formed nanostructure (TD_AT) was measured using DLS. The hydrodynamic size of TD and TD_AT is around 13.06 nm and 30.08 nm respectively (Figure 2 b). The zeta potential of TD which is around −18.84 mV has shifted to +4.5 mV when conjugated with AT. This suggests a greater potential in increasing the uptake of the system due to minimal repulsion due to the negative charge of the outer leaflet of plasma membrane (Figure 2 c, 2 d). The morphology of TD_AT was characterized using Atomic Force Microscopy (AFM) A tetrahedral shape was observed which indicated the formation of AT conjugated tetrahedral framework (Figure 2 f). Since TD’s morphology is well-known and established, we use it (Figure 2 e) to compare with that of our sample (TD_AT) (Figure 2 f). We observed that similar morphology of TD and TD_AT was observed. All the above characterizations collectively prove the successful formation of TD_AT.

### 2.3 In vitro studies

#### 2.3.1 Cellular Uptake studies

The literature shows that TD could enter the cells in 20 minutes [27]. We hypothesized that AT containing TD would be internalized by the cells at a higher rate than TD due to the hydrophobicity of the AT and the inherently high internalisation capacity of TD. Cy3-labelled TD and TD_AT were incubated for 20 minutes in triple negative breast cancer cell lines, MDA-MB-231 and SUM159A. Then the uptake was quantified using confocal laser scanning microscopy Higher the intensity of Cy3, higher the uptake of TD. After quantifying we found that, the uptake of TD_AT was slightly higher in MDA-MB-231 when compared to TD but in SUM159A the uptake of TD and TD_AT was almost similar (Figure 3 a, 3 c).

**Figure 3.**
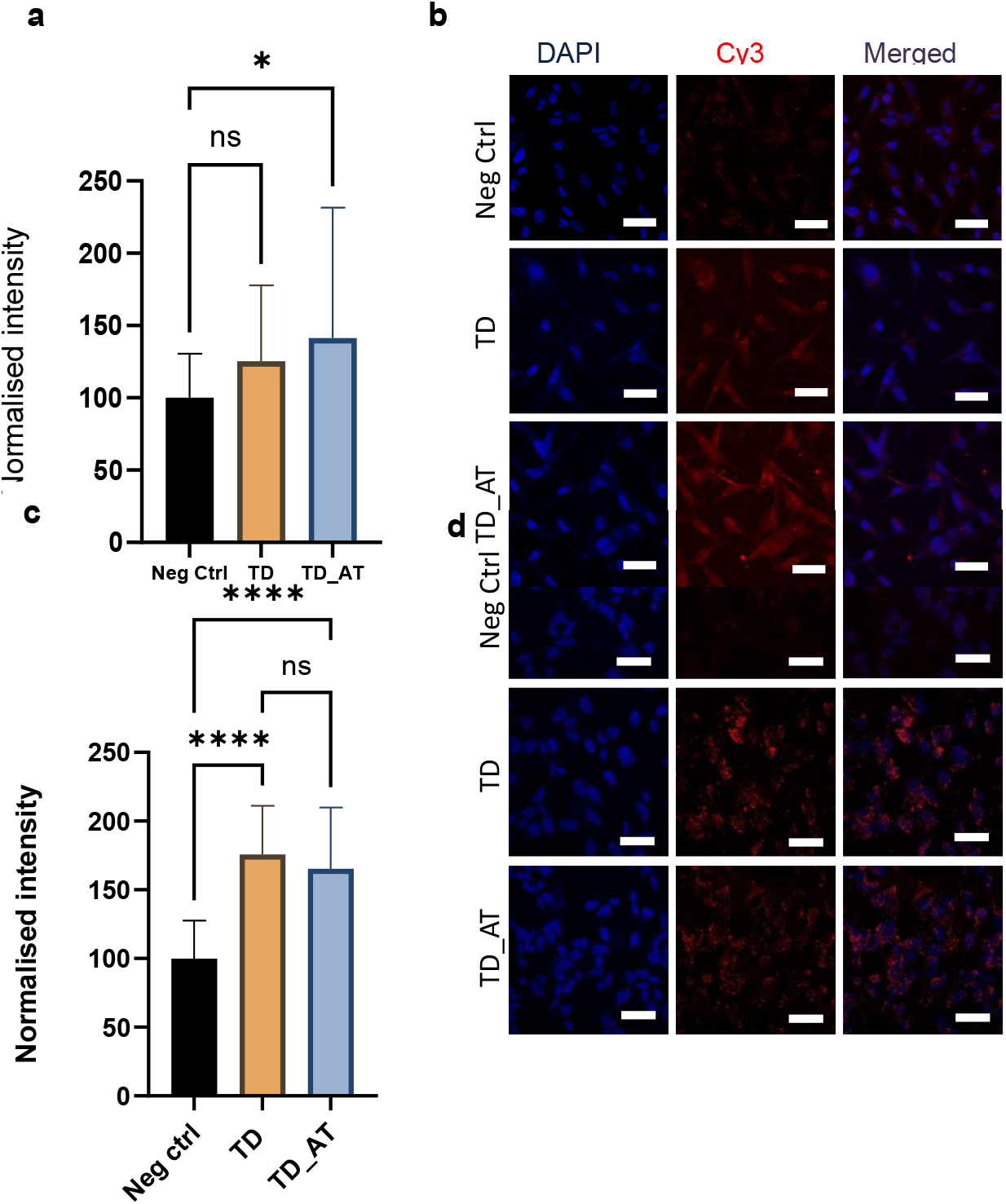
Confocal Images of Cellular Uptake of Cy3 labelled TD and TD_AT. Quantification of fluorescence intensity of the confocal images in the panel (a) MDA-MB-231 (c) SUM159A. Confocal images showing the uptake of TD and TD_AT in (b) MDA-MB-231(d) SUM159A. Blue color shows the DAPI staining the nucleus. Red color indicates the uptake of Cy3 labelled nanostructures. Scale bar is 100 μm

#### 2.3.2 Cytotoxicity studies using MTT assay

Since the knowledge about the cytotoxic effect caused by the system in malignant cells and non-malignant cells are crucial, we studied the cytotoxicity using MTT assay. The experiment was performed in both malignant (MDA-MB-231 & SUM159A) and non-malignant (RPE1) cell lines.

TD itself was known to increase the proliferation and differentiation of the cells due to its antioxidative properties. However, it’s conjugated version (TD_AT) showed significant cytotoxicity in malignant cells but not in non-malignant cells. It was observed that at and beyond 400 nM, TD_AT caused higher cytotoxicity than TD in malignant cell lines MDA-MB-231 and SUM159A. Whereas, in non-malignant cell line (RPE1), TD_AT did not cause any cytotoxicity even at higher concentrations (Ex: 800 nM). (Figure 4). In figure 4d, we could observe that there was nearly 90% viability observed in RPE1 cell lines treated with TD_AT even at concentrations up to 800 nM. On the contrary, in MDA-MB-231 and in SUM159A cells, cell viability started to diminish at 400 nM itself and continued to deteriorate till 800 nM. From this experiment, 400 nM was taken as a concentration of TD_AT which will be used for the further experiments.

**Figure 4.**
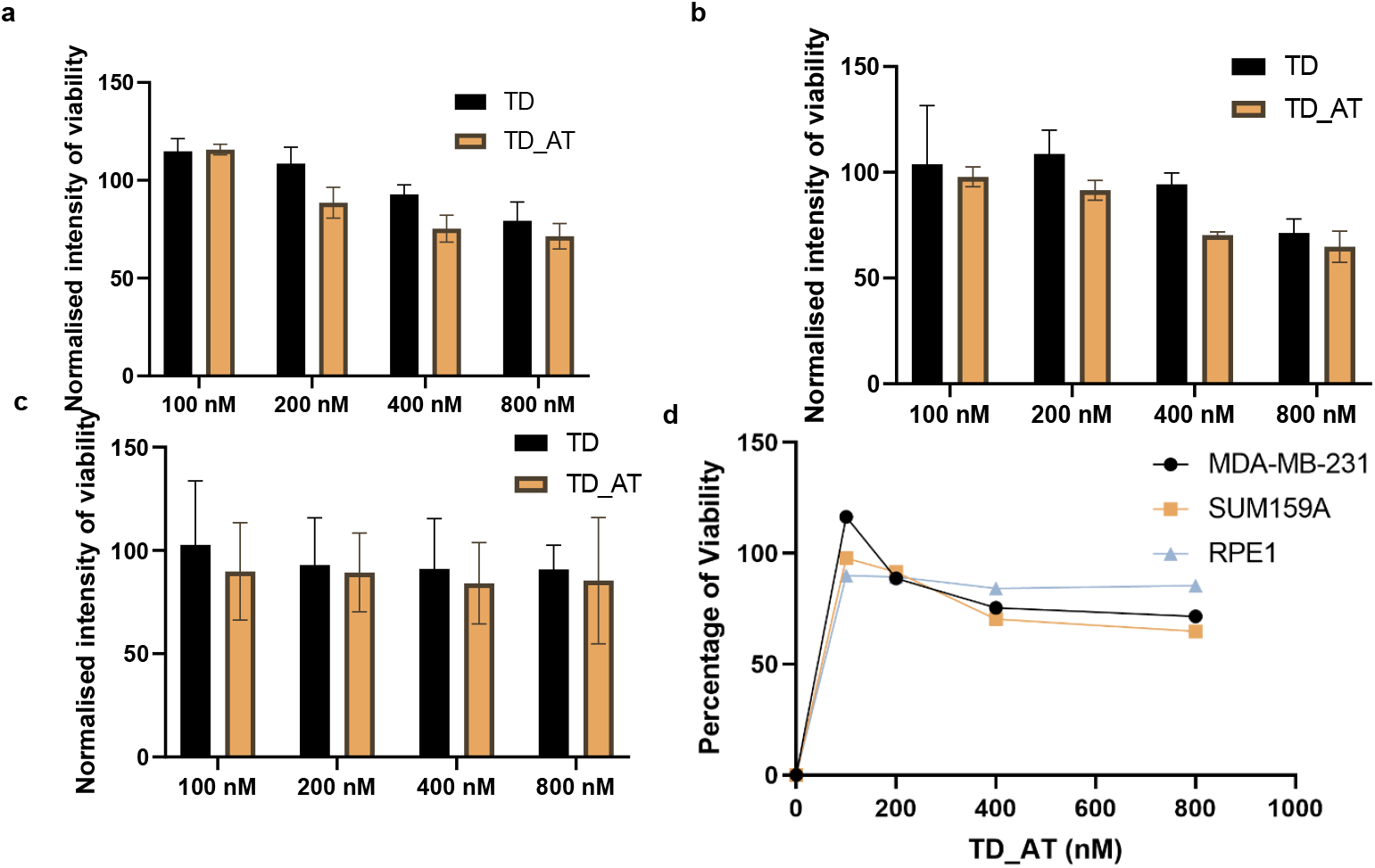
**MTT assay** Plot of normalised intensity of viability indicating the cytotoxicity of TD and TD_AT in (a) MDA-MB-231 (b) SUM159A and (c) RPE1 cell lines. Comparison of all three cell lines was depicted in (d).

**Figure 5.**
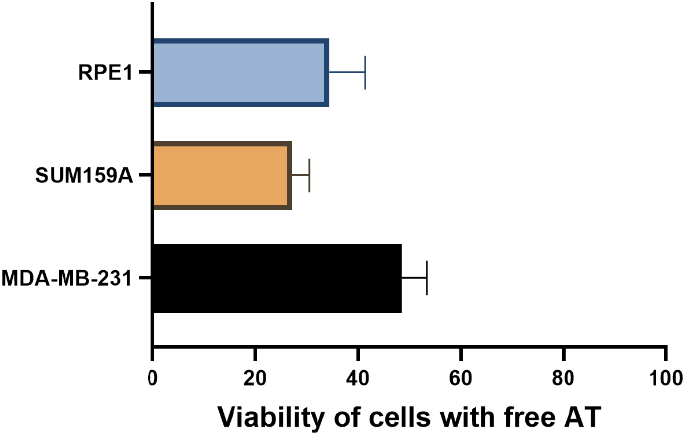
Normalised intensity of viability of cells treated with free AT using MTT assay. Interestingly, from figure 5 we observed that free AT (at a concentration used for coupling) reduced the viability of RPE1 cell lines. From this we can conclude that, the tetrahedron component in TD_AT confers controlled release of the therapeutic molecule (AT) and plays an important role in maintaining the property of selective cytotoxicity of AT and the system itself.

#### 2.3.3 In vitro toxicology studies using LDH assay

By measuring the release of the cytosolic enzyme lactate dehydrogenase (LDH) into the cell culture media, the LDH assay quantifies the damage to cell membranes as a result of cytotoxicity. This release serves as an indication of cell death or damage and is representative of disintegration of the cell membrane. Using a linked enzymatic process, the assay produces NADH when LDH oxidizes lactate to pyruvate. A tetrazolium salt is then reduced by this NADH and transformed into a formazan dye, the quantity of which is directly correlated with LDH activity and cell death. The amount of the formazan product is directly proportional to the amount of LDH released and is the measure of cell death. Therefore, more the formazan, more the LDH released and more the cell damage and cell death. This assay was performed in MDA-MB-231 and RPE1 cell lines.

To validate the results obtained from MTT assay and to confirm the same, we performed LDH assay on both malignant and non-malignant cell lines. From Figure 6a, we could clearly observe that LDH release was observed at comparatively higher amounts in TD_AT than TD in MDA-MB-231 cell line. However, the LDH release was almost same in TD_AT as well as TD in RPE1 cell line (Figure 6 b). As obtained from the MTT assay, the free AT caused significant release of LDH which proves that AT lost its property of selective cytotoxicity at this concentration but retained when conjugated with TD at same concentration. This indicates that TD_AT did not have any effect on the normal cell line (RPE1) but causes significant cytotoxicity in malignant cells (MDA-MB-231). This further validates the findings from the MTT assay.

**Figure 6.**
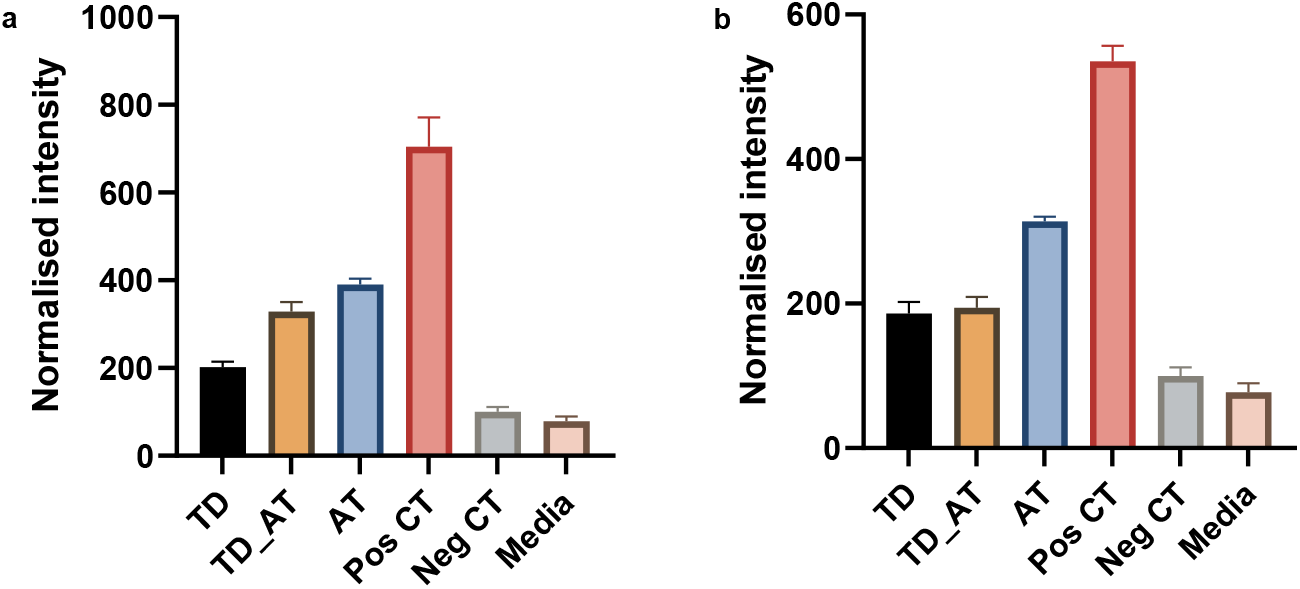
LDH assay performed post-treatment with 400 nM of TD_AT. Normalised intensity of fluorescence determined as a measure of LDH released from (a) MDA-MB-231 and (b) RPE1 cell lines treated with 400 nM of TD and TD_AT.

#### 2.3.4 Intracellular ROS measurement using DCFH-DA

AT when entered the cell in an active state, increases the levels of ROS significantly. To confirm this, we performed DCFH-DA based intracellular ROS measurement assay in which DCFH-DA acts as a probe. DCFH-DA when entered the cell, intracellular esterases deacetylate the probe, which produces DCFH. After which, ROS (Ex: hydrogen peroxide) oxidize DCFH to produce a fluorescent substance dichloro-fluorescein (DCF). DCF’s fluorescence is directly correlated with the quantity of ROS in the cell which is the measure of oxidative stress. We performed this experiment in both malignant (SUM159A) and non-malignant (RPE1) cell line.

It is evident from Figure 7 that higher amounts of ROS was generated for the cells treated with TD_AT when compared to TD. This also suggests that 400 nM concentration of TD_AT is more internalised than 200 nM performed in the previous cellular uptake experiment. Additionally, there was no significant difference in the level of ROS generated in the RPE1 cells treated with TD and TD_AT (Figure 7b). However, free AT generated ROS in both the cases. This further proves the specificity of the system in conferring toxicity only to the malignant cells even when administered the same concentration to the normal cells.

**Figure 7.**
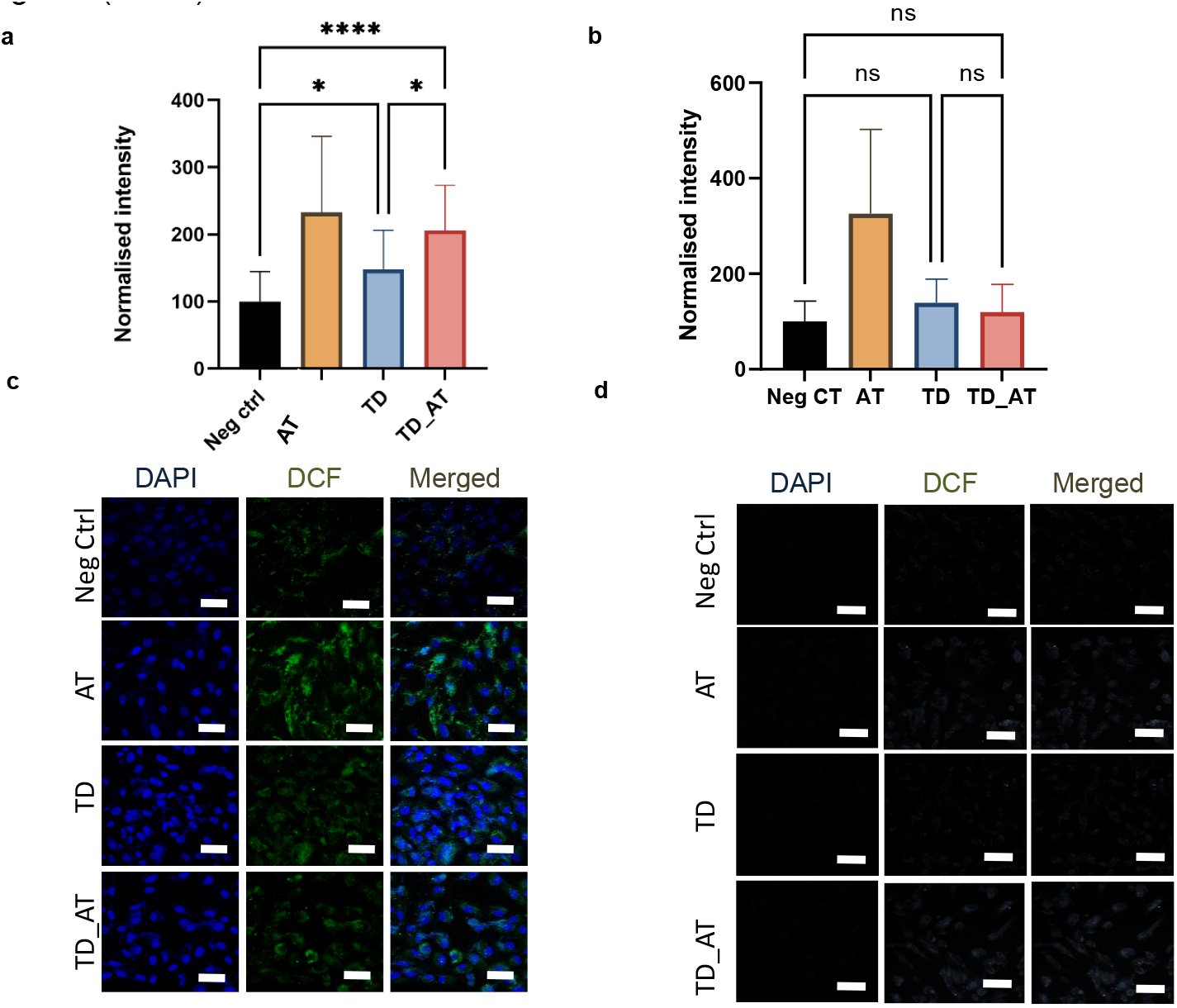
Quantification of intracellular ROS levels post-treatment of 400 nM TD_AT. Quantification of fluorescence intensity of DCFH-DA the confocal images in the panel (a) SUM159A (b) RPE1. Confocal images showing the fluorescence intensity of DCFH-DA in (a) SUM159A (b) RPE1. Blue color shows the DAPI staining the nucleus. Green color indicates the ROS generated. Scale bar is 100 μm

#### 2.3.5 Apoptosis assay

Since earlier experiments confirmed cytotoxicity, we aimed to determine whether it was mediated through apoptosis. We performed apoptosis assay in both adherent malignant (MDA-MB-231) and adherent non-malignant (RPE1) cell lines. Annexin V is an intracellular protein that binds phospholipids and is a member of the annexin family. Phosphatidylserine (PS), which is typically located on the inner side of the plasma membrane, transfers to the outer surface of apoptotic cells. This exposed PS is specifically bound by Annexin V. Conversely, propidium iodide (PI) can penetrate cells with compromised membranes, allowing for the identification of necrotic or late apoptotic cells (positive for both Annexin V and PI) as distinct from early apoptotic cells (Annexin V positive, PI negative).

As we could evidently see from the figure 8c, that increased overall intensity was higher in TD_AT when compared to TD. Comparing the intensity of negative control and test samples, we could arrive at a preliminary confirmation that cell death is mediated by apoptosis. In the case of RPE1 (Figure 8 d), increased intensity was found in free drug (AT) which is close to positive control. Also, the intensity was higher in TD when compared to TD_AT. This again confirms the results from the previous experiments that TD_AT does not cause cytotoxicity to the non-malignant cells and doesn’t lead to apoptosis. Individually quantifying each channel (Annexin V and PI) will give more insights regarding whether the cells are in early apoptotic or late apoptotic stage.

**Figure 8.**
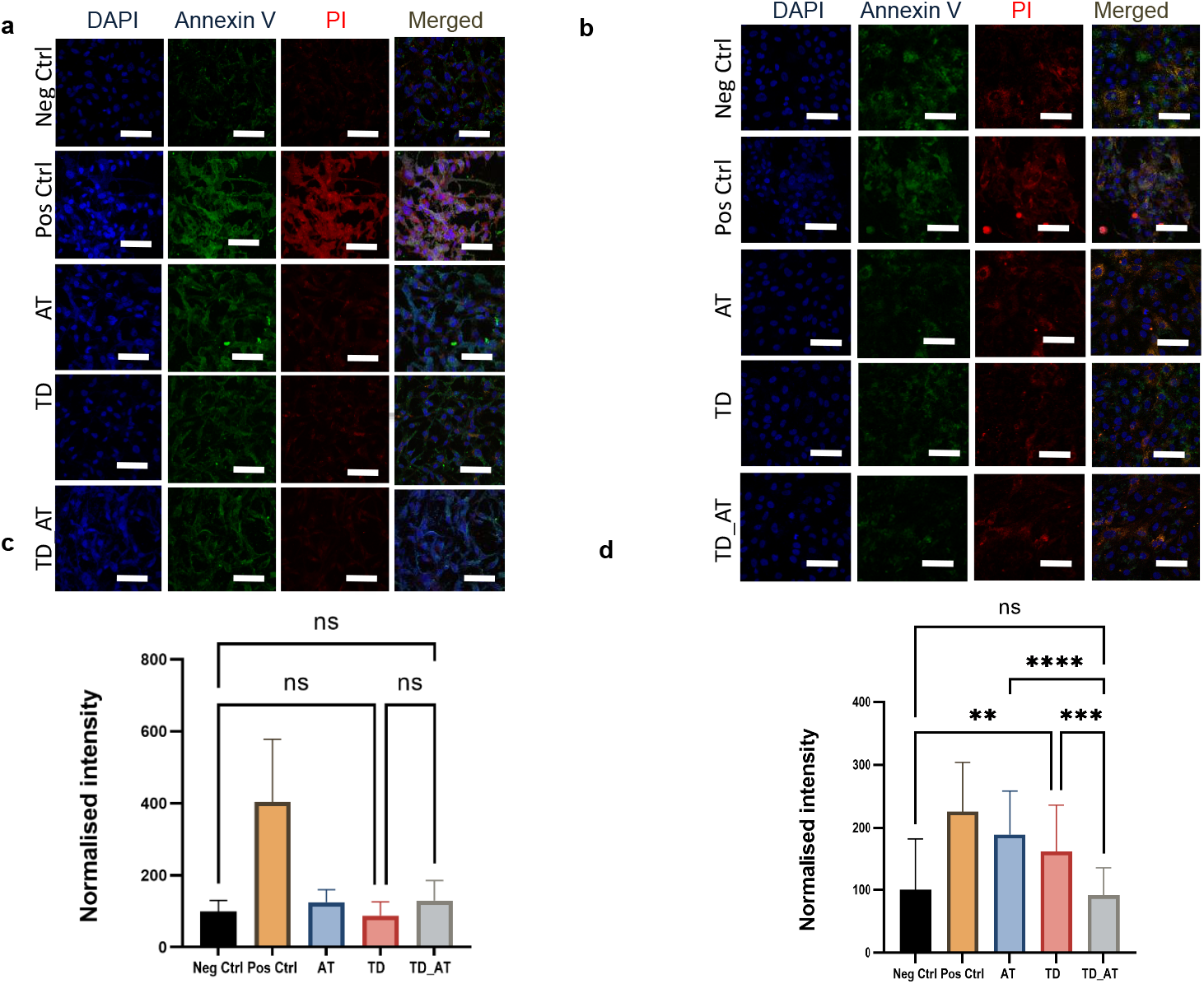
Apoptosis assay using Annexin V and Propidium Iodide on adherent cells. Confocal Microscopy images depicting the apoptosis assay of adherent (a) MDA-MB-231 and (b) RPE1 cells treated with 400 nM of TD_AT and using Annexin V and Propidium Iodide. Quantification of fluorescence intensity of confocal images in the panel (c) MDA-MB-231 (d) RPE1. Scale bar is 100 μm.

## 3. MATERIAL AND METHODS

### 3.1 Materials and Instrumentation

#### 3.1.1 Materials

D-Alpha-tocopherol succinate, MTT reagent (3-(4,5-Dimethylthiazol-2-yl)-2,5-diphenyltetrazolium bromide), the In Vitro Toxicology Assay Kit based on lactate dehydrogenase (LDH), and DNA tetrahedron synthesis primers were obtained from Sigma. Gibco supplied Dulbecco’s Modified Eagle Medium (DMEM), HAM’s F12 medium, fetal bovine serum (FBS), penicillin-streptomycin, trypsin-EDTA (0.25%), and phosphate-buffered saline (PBS). Adherent cell culture dishes were sourced from Thermo Scientific and Tarsons. Chemicals such as paraformaldehyde (PFA), acrylamide:bisacrylamide solution (30%), TEMED, APS, and ethidium bromide were procured from Himedia. Dimethyl sulfoxide (DMSO) was purchased from SRL, while HOBt and EDC were acquired from Avra Chemicals. The Annexin V-FITC/PI apoptosis detection kit was sourced from Elabscience, and desalting columns were purchased from Zeba.

#### 3.1.2 Instruments used

Leica SP-8 confocal microscope, Nikon inverted microscope, Gel electrophoresis apparatus, Mammalian cell culture facilities, Dynamic Light scattering, Atomic force microscope (Bruker), Test Right UV-Visible spectrophotometer, Byonoy Absorbance Microplate Reader, Biorad ChemiDoc MP Imaging System, Malvern analytical Zetasizer Nano ZS instrument.

### 3.2 In vitro cell culture

#### 3.2.1 Cell lines

Human triple negative breast cancer cell lines (MDA-MB-231 & SUM159A) and Human Retinal Pigment Epithelial Cell line (RPE1) were used since they all are epithelial in origin. It would be efficient to compare the effects of malignant epithelial cell lines and non-malignant epithelial cell lines in this study. Also, MDA-MB-231 and SUM159A being highly aggressive and invasive are ideal for studying the response and effect of our systems.

#### 3.2.2 Cell culture

MDA-MB-231 and RPE1 cells were independently maintained in Dulbecco’s Modified Eagle Medium (DMEM) nourished with 10% fetal bovine serum (FBS) and 1% penicillin-streptomycin. SUM159A cells were cultured in HAM’s F12 medium enriched with 10% FBS, insulin, hydrocortisone, HEPES, and 1% penicillin-streptomycin. All cells were incubated in T25 flasks at 37°C in a humidified environment containing 5% CO2. The culture medium was replaced as needed, and cells were passaged upon reaching 80–90% confluency. For passaging, spent medium was removed, and cells were washed with 1X phosphate-buffered saline (PBS) to eliminate residual serum, debris, and trypsin inhibitors that could affect enzymatic action. Cells were then treated with 500 μL of 0.25% Trypsin-EDTA for approximately 4.5 to 5 minutes. Trypsinization was stopped by adding 1 mL of complete medium, and the resulting cell suspension was used for either subculturing into new flasks or seeding into well plates, depending on the experimental needs.

### 3.3 Synthesis of DNA Tetrahedron

#### 3.3.1 Coupling of AT with amino-modified M1 strand

HOBt, AT and EDC were dissolved in DMSO to obtain 50mM, 20 mM and 25mM solutions respectively. HOBt, EDC and AT were added in a ratio of 5:5:1 in DMSO along with 10 μM of amino-modified M1 strand and 20 μM of Triethylamine. The reaction mixture was kept in shaking for 24 hours and in dark conditions for the coupling to take place. After 24 hours, the mixture was then desalted using Zeba spin 7000 MWCO desalting columns. 10 μM solution of AT coupled M1 (M1_AT) strand was obtained and stored at 4°C until further use.

#### 3.3.2 Gel electrophoresis

The coupling was verified using a 10% native PAGE gel. The sample mixture included 5 μL of M1_AT, 3 μL of 1X TAE buffer, and 1.5 μL of 6X gel loading dye. Electrophoresis was carried out at 75 volts for 90 minutes. Following the run, the gel was stained with ethidium bromide for 10 minutes and imaged using the Bio-Rad ChemiDoc MP Imaging System.

#### 3.3.3 UV-Visible Spectrophotometry

To characterize the coupling of AT with amino-modified M1, UV-Visible spectrophotometry was used. The samples were diluted in nuclease-free water in a 1:1 dilution and 300 μL was analysed using Test Right UV-Visible spectroscopy. Characteristic peak of DNA at 260 nm and of AT around 250 nm was used to assess the shift in peak due to coupling.

#### 3.3.4 One-pot synthesis of DNA Tetrahedron

100 μM stocks of single stranded oligonucleotides M1, M2, M3, M4, Cy3 modified M4 (primers) were reconstituted using nuclease free water accordingly by subjecting to 350 rpm at 70°C for 1.5 hours. From this, 10 μM solution was diluted using nuclease free water or 1X PBS according to the requirement of the experiment. M1, M2, M3 and M4 oligonucleotides were taken in equimolar concentrations for the formation of DNA TD. 2mM MgCl2 was added to reduce the intramolecular repulsion between the oligonucleotides and to increase the rigidity and stability of the duplex. Thermal annealing was performed in a thermocycler, beginning at 95°C, holding for 30 minutes and gradually cooling down to 4°C in 5°C decrements, with each step held for 15 minutes. The resulting DNA tetrahedron (TD) was obtained at a final concentration of 2.5 μM and stored at 4°C for subsequent use. For the synthesis of TD_AT, same procedure was performed with M1_AT in place of M1.

**Table 1.**
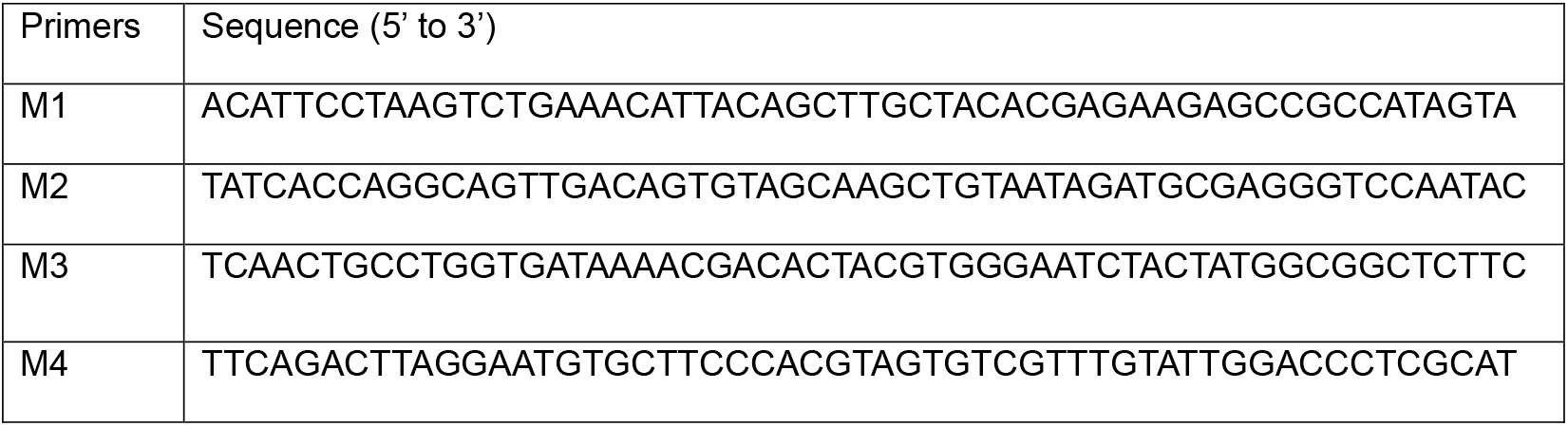
Sequences of oligonucleotides used for TD synthesis.

#### 3.3.5 Electrophoretic mobility shift assay (EMSA)

Electrophoretic mobility shift assay (EMSA) was conducted using a 10% native PAGE gel to verify the formation of the higher-order structure (TD_AT). The reaction mixture comprised 5 μL of TD, 3 μL of 1× TAE buffer, and 1.5 μL of 6× loading dye. Electrophoresis was carried out at 90 volts for 90 minutes. After the run, the gel was stained with ethidium bromide for 10 minutes and imaged using the Bio-Rad ChemiDoc MP Imaging System.

#### 3.3.6 Dynamic Light Scattering (DLS)

To measure the hydrodynamic size of the formed nanostructure, DLS was performed. 100 μL of 2.5 μM sample was diluted in 1:10 dilution, centrifuged at 10,000 rpm for 10 minutes. 950 μL of supernatant was collected, made up to 1 ml using nuclease-free water and used for analysing the hydrodynamic radius using Malvern analytical Zetasizer Nano ZS instrument. The readings were taken in triplicates for accuracy.

#### 3.3.7 Atomic Force Microscopy (AFM)

Atomic Force Microscopy (AFM) was employed to characterize the morphology of the nanostructure. The sample was diluted at a 1:2 ratio, applied onto a freshly cleaved mica surface, and vacuum-dried for a minimum of 24 hours. Imaging was performed using a Bruker AFM in tapping mode with a cantilever tip. The resulting images were processed and analysed using JPK software.

### 3.4 In vitro mammalian cell culture studies

#### 3.4.1 Cellular Uptake studies

Cellular uptake experiments were conducted using two malignant cell lines, MDA-MB-231 and SUM159A. Cells from T25 flasks at 80–90% confluency were seeded onto 12 mm coverslips placed in 4-well plates at a seeding density of 0.05 × 10^6^ cells per well. After allowing the cells to adhere for 24 hours in an incubator, and once they reached approximately 80% confluency, the spent medium was removed to initiate the experiment. Cells were washed with 1X PBS and incubated in serum-free medium for 20 minutes to assess stress response. Upon confirming that the cells are healthy using a microscope, the medium was replaced with serum-free medium containing 200 nM of Cy3-labeled TD and TD_AT, followed by a 20-minute incubation at 37°C. Post incubation, cells were washed twice with 1X PBS to eliminate unbound nanostructures. Fixation was performed using 4% paraformaldehyde at 37°C for 15 minutes. Cells were then washed three times with 1X PBS to remove residual fixative. Coverslips were mounted onto glass slides using DAPI-containing Mowiol and left to dry overnight at 4°C. Imaging was carried out using a Leica SP8 confocal microscope with a 63X oil immersion objective and appropriate laser settings. The acquired images were analysed using ImageJ software.

#### 3.4.2 Cell viability studies using MTT assay

Cell viability assays were performed on both malignant (MDA-MB-231, SUM159A) and non-malignant (RPE1) cell lines. Cells from T25 flasks at approximately 85% confluency were seeded into 96-well plates at a density of 20,000 cells per well and allowed to adhere for 24 hours at 37°C. Once the cells reached around 80% confluency, they were treated with varying concentrations (100, 200, 400, and 800 nM) of TD and TD_AT in complete medium for 24 hours. Following treatment, the media was discarded, and cells were washed with 1X PBS. An MTT solution (0.5 mg/mL of 3-(4,5-Dimethylthiazol-2-yl)-2,5-diphenyl tetrazolium bromide) was added to each well, followed by a 4-hour incubation at 37°C. Post incubation, the MTT solution was removed, and 100 μL of DMSO was added to each well and incubated in the dark for 10 minutes to solubilize the formazan crystals. Absorbance was measured at 562 nm using the Byonoy Absorbance Microplate Reader. The resulting data were plotted and analysed using GraphPad Prism software.

#### 3.4.3 Intracellular ROS measurement using DCFH-DA

Intracellular reactive oxygen species (ROS) production was assessed using DCFH-DA (Dichloro-dihydro-fluorescein diacetate) in MDA-MB-231, SUM159A, and RPE1 cell lines. Cells from T25 flasks at 80-90% confluency were seeded onto 12 mm coverslips in 4-well plates at a density of 0.05 × 10^6^ cells per well and allowed to adhere for 24 hours at 37°C. Once cells reached 80% confluency, the spent medium was discarded, and cells were washed twice with 1X PBS. They were then incubated with 400 nM TD_AT in serum-free medium for 2 hours at 37°C. After incubation, the cells were washed again with 1X PBS and incubated with 10 μM DCFH-DA for 20 minutes at 37°C. Following this, the cells were washed twice with 1X PBS and fixed with 4% paraformaldehyde for 15 minutes at 37°C. The fixed cells were washed three times with PBS and mounted onto glass slides using DAPI-containing Mowiol, then left to dry overnight at 4°C. Imaging was performed using a Leica SP8 confocal microscope with a 63X oil immersion objective and appropriate laser settings. The acquired images were processed and analysed using ImageJ software.

#### 3.4.4 In vitro toxicology studies using Lactic dehydrogenase assay (LDH assay)

In vitro toxicology studies were performed on both malignant (MDA-MB-231) and non-malignant (RPE1) cell lines using an LDH assay kit to measure LDH release into the culture medium. A 1X LDH assay cofactor solution was prepared by dissolving lyophilized cofactor in 25 mL of autoclaved Type I water, and aliquots were stored at −20°C. The LDH assay mixture was freshly prepared by combining equal volumes of LDH substrate solution, LDH assay dye solution, and 1X LDH cofactor solution. Cells were seeded into a 96-well plate at a density of 20,000 cells per well from T25 flasks at 85% confluency and allowed to adhere for 24 hours at 37°C. Once they reached 80% confluency, cells were treated with 400 nM of TD or TD_AT in complete media for 24 hours. After treatment, the spent media was collected and centrifuged at 250g for 10 minutes to remove debris and any remaining cells. The supernatant was transferred to a clean 96-well plate for enzymatic analysis. An equal volume of LDH assay mixture was added, and the plate was incubated at room temperature for 20– 30 minutes. To stop the reaction, 1/10th volume of 1N hydrochloric acid was added. Absorbance was measured at 490 nm using the Byonoy Absorbance Microplate Reader. The data were plotted and analysed using GraphPad Prism.

#### 3.4.5 Apoptosis assay using adherent cells

Apoptosis was assessed using the Annexin V-FITC/PI apoptosis kit in both malignant (MDA-MB-231) and non-malignant (RPE1) cell lines. Cells from T25 flasks at 80-90% confluency were seeded onto 12 mm coverslips in 4-well plates at a density of 0.05 × 10^6^ cells per well and allowed to adhere for 24 hours at 37°C. Once cells reached 80% confluency, the spent medium was removed, and cells were washed with 1X PBS followed by 1X Annexin V binding buffer. Cells were then incubated with Annexin V-FITC and Propidium Iodide (2.5 μL each) in 400 μL of binding buffer for 20 minutes in the dark at room temperature. After incubation, cells were gently washed with 1X PBS and fixed with 2% paraformaldehyde for 15 minutes at 37°C. The fixed cells were washed three times with PBS and mounted onto glass slides using DAPI-containing Mowiol, then left to dry overnight at 4°C. Imaging was performed using a Leica SP8 confocal microscope with a 63X oil immersion objective and appropriate laser settings. The images were processed and analysed using ImageJ software.

#### 3.4.6 Confocal Microscopy

Fixed-cell imaging was performed using a Leica TCS SP8 confocal laser scanning microscope with a 63× oil immersion objective and a resolution of 512 × 512 pixels. The pinhole was set to 1 Airy unit, and imaging parameters including bit depth, laser power, and detector gain were kept constant across all experimental conditions within a given study. Fluorophores were excited using appropriate laser lines: DAPI at 405 nm with a Diode 405 laser; DCFDA and Annexin V-FITC at 488 nm with an Argon laser; Cy3 and Propidium Iodide at 561 nm with a DPSS laser. Emission detection windows were configured according to the specific emission spectra of each fluorophore. Z-stack imaging was performed with system-optimized step sizes. For experiments containing multiple fluorophores, sequential scanning was employed to minimize spectral overlap and crosstalk.

#### 3.4.7 Image Processing and Statistical Analysis

Image analysis was conducted using Fiji (ImageJ). Z-stack images were converted into two-dimensional representations using the Z-projection function, specifically employing the maximum intensity projection method. Background correction was performed by subtracting the average background intensity from each image. Following this, quantitative measurements of integrated density and raw integrated density were obtained. Data normalization and statistical analyses were carried out using GraphPad Prism (version 9.0). For normalization, a baseline value of zero was set as 0% across all datasets, while the average integrated density of the control group was defined as 100%. Statistical comparisons between two groups were performed using unpaired, two-tailed t-tests, while comparisons involving more than two groups were analysed by one-way. All statistical tests were conducted under the assumption of normally distributed data.

## 4. Conclusions and Future Directions

Conventional cancer therapies suffer from disadvantages like poor bioavailability, low solubility and especially, off-target effects. Therapeutics like monoclonal antibodies, antibody-drug conjugates and targeted nanoparticles being advantageous in delivering to the targeted area with high specificity poses a threat of burst release, cumbersome and expensive synthesis process and larger size. This shifts the focus to a budding arena, DNA nanotechnology in which DNA nanocages were well established for their applications in drug delivery, bioimaging, biosensing, nanomedicine, etc due to their smaller size, biocompatibility, programmability and ease of use. This study investigates the potential of DNA tetrahedron (TD), a DNA nanocage, in delivering alpha-tocopherol succinate (AT), an underrated molecule known for its unique property of being selectively cytotoxic to malignant cells when used at appropriate concentrations. On investigating whether the components (AT and TD) synergistically help each other in overcoming their drawbacks or not, in this study we sought to fabricate a novel system containing both TD and AT.

In order to overcome the disadvantages caused due to non-covalent binding, AT was conjugated to DNA TD (TD_AT) via HOBt-EDC chemistry which was characterized using various techniques namely electrophoretic mobility shift assay, UV-Visible spectrophotometry, Atomic force microscopy and Dynamic Light Scattering. Invitro studies focussing on cellular uptake of the system (TD_AT) yielded increased uptake of TD_AT in MDA-MB-231 cell line but not in SUM159A cell line. This might be due to the variability in the membrane components (SUM159A being positive for N-cadherin and E-cadherin unlike MDA-MB-231. Cytotoxicity studies using MTT assay proved the selective cytotoxic property of the system to malignant cells. Toxicology studies using LDH release assay and intracellular ROS assay using DCFH-DA also confirmed the outcome of MTT assay that TD_AT treated malignant cells released more LDH and generated more ROS when compared to TD. In normal cells (RPE1), the toxicity and ROS generation was less when compared to the TD treated cells. Interestingly, free AT treated normal cells showed toxicity to the compound whereas TD_AT treated cells bypassed this with no toxicity. This showed that TD played a significant role in retaining the unique property of AT. Apoptosis assay further confirmed the results which indicates a preliminary conformation that cell death is mediated by apoptosis. Flow cytometry-based apoptosis studies might yield a better conclusion about the stages of apoptosis. Conjugating evert strand of TD with AT might increase the efficiency of the system. Studies on mitochondrial stability might yield useful insights about the pathway and mechanism of cytotoxicity caused

## Acknowledgements

We sincere thanks all the members of DB Lab at IITGN for useful discussions and inputs. We thank Central Instrumentation Facility at IITGN for the infrastructure. DB thanks ANRF-CRG, GSBTM and MoES-STARS for research funding.

The authors declare no conflict of interest.

## Notes

### Competing Interest Statement

The authors have declared no competing interest.

